# Soft-lithographically defined template for arbitrarily patterned acoustic bioassembly

**DOI:** 10.1101/2024.04.07.588443

**Authors:** Sihan Chen, Jibo Wang, Shanqing Jiang, Yuhang Fan, Yang Zhao, Wen Zhao, Zian Wan, Qin Zhou, Yun Chen, Pu Chen

## Abstract

Acoustic bioassembly is recently regarded as a highly efficient biofabrication tool to generate functional tissue mimics. Despite their capacity of directly patterning live cells with close intercellular proximity, most acoustic bioassembly techniques are currently limited to generate some specific simple types of periodic and symmetric patterns, which represents an urgent challenge to emulate geometrically complex cytoarchitecture in human tissue. To address this challenge, we herein demonstrate a soft-lithographically defined acoustic bioassembly (SLAB) technique that enables to assemble live cells into geometrically defined arbitrary multicellular structures. Particularly, we employed a widely accessible soft lithography technique to fabricate a PDMS construct that works as an amplitude modulation template to define the pressure distribution of near-field acoustic waves. We found that zero pressure areas of the near-field acoustic waves at the PDMS surface distribute above the air-filling regions of the PDMS construct when both the PDMS top layer and air layer are approximately one-tenth of the acoustic wavelength. Using this technique, bioparticles can be assembled into symmetrical or asymmetrical patterns. Specifically, we have demonstrated the SLAB of endothelial spheroids and hepatic cells into liver tissue mimics (LTMs). The functional analysis further indicates that the formed LTMs displayed liver-specific functions, including albumin secretion, urea synthesis, glucose metabolism, and lipid storage. We expect this SLAB technique will be broadly used to construct complex functional tissues for tissue engineering and regenerative medicine.

## 1. Introduction

Advanced biofabrication is an essential basis for tissue engineering and regenerative medicine, and it aims to generate microscale human-relevant tissue mimics for basic medical research as well as macroscale tissue substitutes for clinical regenerative therapy^1–3^. Specifically, bioprinting is a widely-adopted technical route of biofabrication. Traditional bioprinting techniques, such as inkjet printing and extrusion printing, enable to directly print biomaterials to form arbitrary complex 3D biomaterial construct^4^. Despite the structure of printed biomaterials being arbitrarily defined, live cells in the biomaterial are randomly and loosely distributed and cannot form close intercellular contact^5, 6^. As an alternative approach to bioprinting, bioassembly has recently been regarded as a critical technical route of biofabrication by the International Society of Biofabrication (ISBF)^7, 8^. Bioassembly employs interactions of the physical fields and biological elements (i.e., cells, spheroids, and organoids) to manipulate live cells directly. Cells in the physical field self-assemble into closely packed multicellular structures at the locations where force potential is minimized. The close intercellular proximity facilitates contact-dependent cell communications via gap and tight junctions, ultimately enhancing cell coordination, polarization, and maturation of tissue-specific functions^9, 10^.

Currently, bioassembly has been exploited to construct a variety of tissue mimics using various physical fields, including optical, electrical, magnetic, and acoustic fields^11–16^. Particularly, acoustic bioassembly techniques hold a better promise to become an efficient biofabrication tool than the other bioassembly techniques. Acoustic bioassembly displays several advantages, such as flexibility in tuning assembly patterns, scalability in producing macroscale structure, and biocompatibility with primary cells and hPSC-derived adult cells^17–19^. Specifically, acoustic standing waves, such as surface acoustic waves and bulk acoustic waves, have been employed to generate multicellular tissue constructs with simple types of node or antinode patterns, such as cluster array, line array, and grid^20, 21^. Notably, Faraday waves, a standing wave induced by vertical mechanical vibration, enable to generate more complex periodic and symmetric patterns with symmetrical boundaries or asymmetrical patterns with non-symmetrical boundaries^22, 23^. However, acoustic bioassembly patterns generated by standing waves are confined to the regions of either acoustic pressure nodes or antinodes and cannot be arbitrarily defined. Therefore, these standing wave-based acoustic bioassembly techniques could not fully fill the tasks of advanced biofabrication, which presents a critical challenge to acoustic bioassembly.

To address this challenge, a few novel acoustic bioassembly techniques have been recently demonstrated using traveling wave-based acoustic techniques^24, 25^. Specifically, acoustic holography (AH) enables to generate arbitrarily spatial distribution of bioparticles on a specific assembly plane^26^. The AH platform usually contains an acoustic transducer and a custom-designed acoustic holographic kinoform (*i.e*., metamaterial) that spatially modulates the phase of the incident acoustic plane wave from the transducer. Thus, a predesigned arbitrary acoustic pressure distribution can be achieved for bioparticle assembly. As an alternative configuration to the acoustic holographic kinoform, a phased array composed of multiple acoustic transducers can be used to modulate the wavefront of propagating waves. Whereas, the AH is limited to manipulate relatively large and low-density particles due to the diffraction limit of acoustic holographic kinoform. Recently, acoustofluidic holography (AFH) exceeds the particle manipulation limit of AH and enables to manipulate individual cells and even nanoscale particles at the liquid-air interface. The AFH explored kinoform modulated acoustic radiation force and acoustofluidic flow-induced hydrodynamic drag force^27^. While these phase holography techniques have demonstrated great potential in constructing the desired cytoarchitecture, numerical design and fabrication of acoustic kinoform are relatively highly complex.

Here, we demonstrate a soft-lithographically defined acoustic bioassembly (SLAB) technique that does not rely on complex devices and algorithms such as programmed metasurface to form arbitrarily geometric multicellular structures. Specifically, we utilize soft lithography, a widely accessible microfabrication technique, to generate a PDMS construct based amplitude modulation template (AMT) that determines the pressure of near-field acoustic waves. We numerically and experimentally investigate the effects of PDMS top layer, air layer, and bioparticle size on acoustic bioassembly patterns. As a proof of concept, we utilized this technique to fabricate liver tissue mimics (LTMs), and performed functional assessments on the formed LTMs, including albumin secretion, urea synthesis, glycogen storage, and lipid accumulation.

## 2. Materials and methods

### 2.1 Fabrication of acoustic modulation template

We fabricated an AMT using polydimethylsiloxane (PDMS) soft-lithography. Specifically, the lithographic photomask was designed to match the target pattern of bioassembly using AutoCAD 2022 (*Autodesk, USA*). The positive mold of the SU-8 photoresist (*Microchem, Westborough, MA*) was fabricated using a photomask and a mask aligner (*Xueze, JKG-2A*). PDMS (*Dow Corning, Sylgard 184*) construct was fabricated using the replica molding. A PDMS thin film with a thickness of 50 μm was fabricated by spin-coating of PDMS prepolymer and subsequent baking. The AMT was generated by bonding the microfabricated PDMS construct to the petri dish via oxygen plasma treatment, as shown in Fig. 1A.

**Figure 1:**
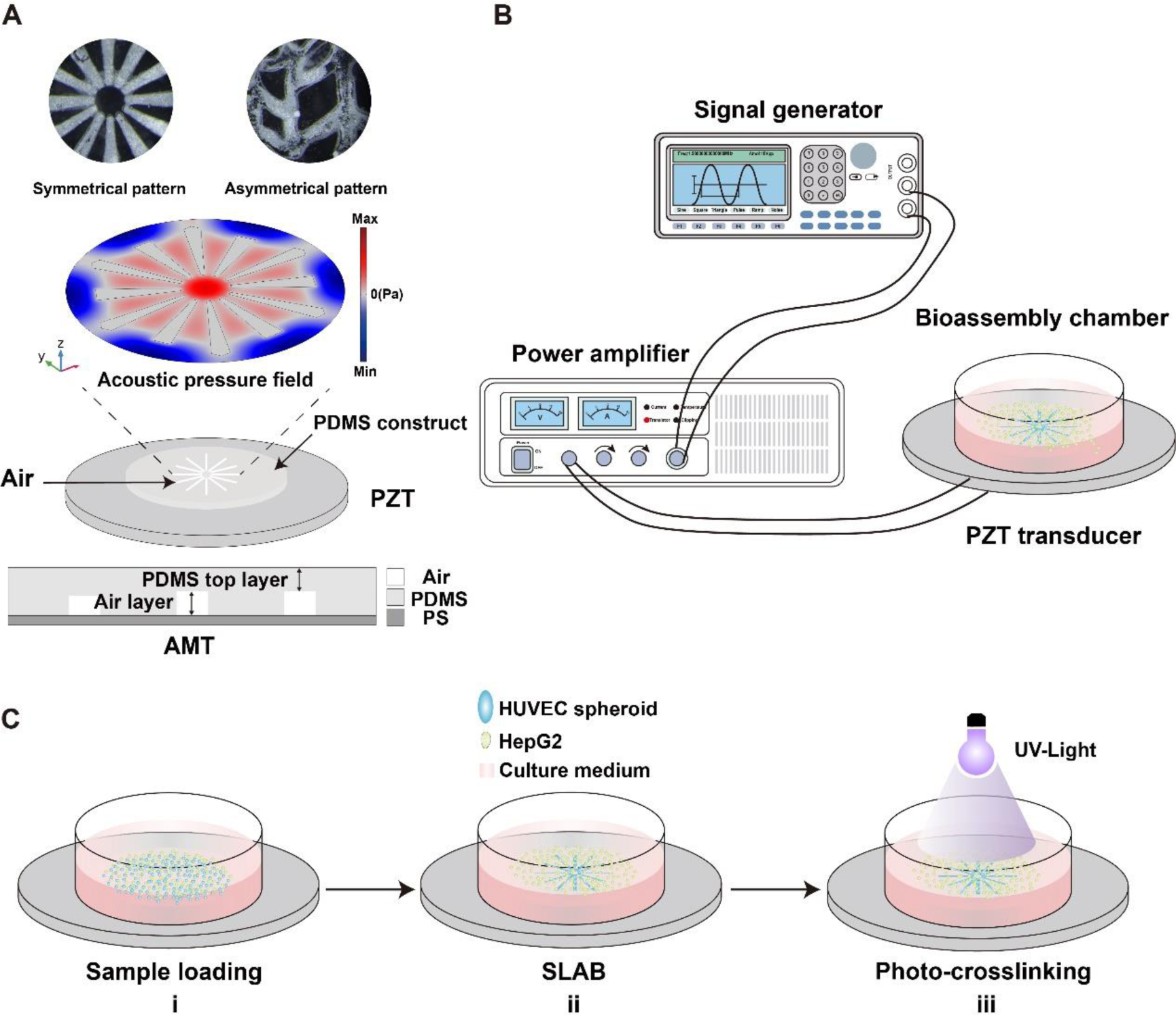
Scheme of SLAB. (A) Working principle of SLAB. Bulk acoustic waves are generated by PZT and then modulated by an AMT. The acoustic pressure node formed on the air-filling regions of the PDMS construct. Microparticles were assembled on the acoustic pressure nodes. (B) Experimental apparatus of SLAB includes signal generator, power amplifier, assembly chamber and PZT transducer. (C) Schematic demonstration of the procedure for constructing a LTMs by SLAB.

### 2.2 Experimental setup

The experimental device consisted of a piezoelectric transducer (*Fuji Ceramics corporation*), a function signal generator (*Tektronix, AFG3052C*), a power amplifier (*Falco system, WMA-300*), and a bioassembly chamber. A piezoelectric transducer (PZT) was composed of piezoelectric ceramic embedded on a polymethylmethacrylate (PMMA) sheet (dimension, φ25 mm × 1 mm; resonant frequency, 1.9 MHz). The function signal generator provided a 1.9 MHz AC sine electrical signal to the power amplifier. The PZT was driven by the amplified signals and generated bulk acoustic waves. The bioassembly chamber was engineered from a commercialized 3.5-cm cell culture dish (*Corning, NY 14831*). The AMT was bonded on the surface of the cell culture dish. The bioassembly chamber was mounted on the PZT through an ultrasonic coupling agent (*Cofoe Corporation*), as shown in Fig. 1B.

### 2.3. Numerical Simulation

We performed numerical simulation to investigate acoustic pressure field, acoustic stream velocity, and their effect on particle motion. Acoustic radiation force (Frad) acting on the small particles was calculated using the Gor’kov theory^28, 29^. In this theory, *F^rad^* is calculated by

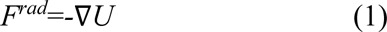

where U is the Gor’kov potential, given by

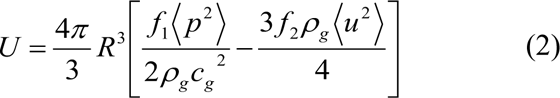

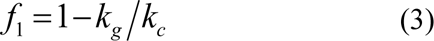

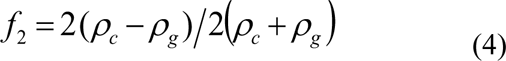

where *p* and *u* are the acoustic pressure and velocity, respectively. *ρ_g_* and *ρ_c_* are the density of the fluid and cell, *c_g_* is the speed of sound in fluid, *k_g_* and *k_c_* are bulk modulus of the fluid and cell, respectively. The Gor’kov theory is valid when the sphere radius is much smaller than the wavelength *λ*.

All the numerical simulations were conducted using 3D finite element method with COMSOL Multiphysics 6.0 (*COMSOL Inc., Burlington, MA, USA*). The “Pressure Acoustics, Frequency Domain” module was used to model acoustic wave propagation and obtain the fluid domain’s pressure field distribution. The “Solid Mechanics” module was used to define the boundary conditions of piezoelectric ceramic, and the “Electrostatic” module defined the input signals on piezoelectric ceramic. The “Particle Tracing” module was utilized to track particle movements within the prepolymer solution under the acoustic radiation force. The “Electric Potential” condition was applied to the domain of “Electrostatic” physics. The boundary conditions on the solid-liquid interface were set to “Acoustic structure interaction”. The simulations were solved via a “Frequency Domain” solver. The material properties used in the calculation are given in Table 1.

### 2.4 Microparticle assembly

Four types of polystyrene (PS) microparticles (diameters of 10, 50, 100, and 200 μm) were used in the microparticle assembly. The microparticle suspension was prepared by suspending microparticles in the phosphate buffer solution (PBS) in the bioassembly chamber. The microparticle suspension was assembled under various concentrations (1, 1.5, and 2 mg/mL) and assembly times (0, 10, and 60 s) by applying a 1.9 MHz electrical signal with an amplitude of 50 Vpp to the PZT. Images was acquired using a stereomicroscope (*Sunny Optical Technology, SZX12*).

### 2.5 Cell culture

Green fluorescent protein (GFP) labeled endothelial cells were cultured in endothelial cell culture medium (ECM) (*ScienCell, CA, USA*) with 5% (v/v) fetal bovine serum (FBS), 1% (v/v) endothelial cell growth supplement (ECGS) and 1% (v/v) penicillin/streptomycin. After cells reached confluent, endothelial cells were digested with 0.05% trypsin EDTA (*Gibco, CA, USA*) and then used to generate endothelial spheroids.

HepG2 cells were cultured in Dulbecco’s modified eagle medium (DMEM) (*Gibco, CA, USA*) with 5% (v/v) FBS and 1% (v/v) penicillin/streptomycin. After reaching confluence, HepG2 cells were treated with 0.05% trypsin EDTA to produce single-cell suspension.

### 2.6 Endothelial spheroid preparation

Endothelial spheroids were generated using a homemade microcavity array chip^30^. Specifically, the layout of the microcavity arrays was initially designed using AutoCAD, and a PMMA female mold was fabricated on a PMMA plate (*CREROMEM*) using a precision micro-milling machine (*JINGYAN, E3020*) equipped with a parabolic spiral-flute bit (*WeiTol*). Then, the PDMS male mold was fabricated by pouring PDMS prepolymer (ratio 1:10, cross-linking agent: elastomer) to the PMMA female mold and solidifying the PDMS prepolymer at 80℃ for 2 h. The released PDMS mold was then immersed in a preheated agarose solution (2% w/v). The agarose-based microcavity arrays were formed by solidifying the agarose solution at 4℃ for 20 min and released from the PDMS mold. The microcavity arrays were removed from the agarose construct using a round punch and gently placed into a 24-well plate. The microcavity arrays in the plate were sterilized thoroughly in the purified water with 1% (w/v) penicillin-streptomycin and then under ultraviolet light for more than 30 min. After 1×10^6^/well of endothelial cells were inoculated into the agarose array, the 24-well plate was placed on a 75-rpm orbital shaker for dynamic culture with ECM, maintained for four days, and replaced with the culture medium every other day.

### 2.7 Acoustic bioassembly of liver tissue mimics

The mixture comprising hepatic cells (1×10^7^/mL) and endothelial spheroids (1.5×10^4^/mL) was prepared in a 2.5 mg/mL GelMA prepolymer solution (*EFL, GM-30*). Then, 20 μL of the mixture was added into the bioassembly chamber before applying acoustic fields. Once RF signals at 1.9 MHz were applied, the cells were assembled on the pressure nodes and formed the target pattern. The assembled multicellular construct was immobilized with UV crosslinking (Fig. 1C) and was cultured at 5% CO_2_ and 37°C in a humidified incubator using a 1:1 mixture of DMEM and ECM for 7 days. The culture medium was refreshed once every 2 days.

### 2.8 Cell viability assay

The cell viability of hepatic and endothelial cells was examined by Hoechst 33342/PI staining. Tissue constructs were washed with 1× PBS 3 times and incubated for 30 min at 37°C in 1 mL staining solution containing 0.1% (v/v) Hoechst 33342 solution (*Cell Signaling Technology, MA, USA*) and 0.1% (v/v) PI solution (*Beyotime Biotechnology, Shanghai, China*). After that, tissue constructs were imaged under an inverted fluorescence microscope (*IX83, Olympus, Japan*) with a 10× objective. The percentage of area positively stained for PI was determined using ImageJ software (*NIH, Bethesda, MD, USA*) and was relative to the total area positively stained for Hoechst 33342. Image analysis was conducted using ImageJ to quantify cell viability rate.

### 2.9 Cell imaging

Vital staining was used to visualize the spatial distribution of heterocellular architecture in the LTMs. According to the manufacture protocol, hepatic cells were stained with CellTracker fluorescent probes (*Life Technologies*), a red-color fluorescent dye. HUVECs were stably expressing GFP. The LTMs were imaged under an inverted fluorescence microscope with a 4× objective (*IX83, Olympus, Japan*).

### 2.10 Liver functional assays

Human albumin secretion of the LTMs was measured using a human albumin ELISA quantitation set (*Bethyl Laboratory, TX, USA*). Urea secretion of the LTMs was determined using a urea assay kit (*BioAssay Systems, CA, USA*). The tissue culture supernatants were collected on days 2, 4, and 6 for the secretion assays. The glycogen synthesis function of the LTMs was assessed using Periodate Schiff (PAS) (*Leagene, Beijing, China*) staining. Lipid storage function was assessed using Nile red staining (*Sigma, MO, USA*).

### 2.11 Statistical analysis

Data presented were quantified from at least three independent experiments. One-way analysis of variance (ANOVA) with Student’s t-test was used to verify statistically significant differences among experiment groups (ns depicts P > 0.05). All values were presented as mean ± standard error of the mean (s.e.m). All collected data were analyzed using the statistical software GraphPad Prism 8.0.0 (*GraphPad Software, San Diego, CA, USA*).

## 3. Results

### 3.1. Numerical simulation of acoustic waves

We investigated the dimension effects of the AMT on acoustic bioassembly patterns using numerical simulation. Specifically, an AMT was designed with an array of circular-shaped air-filling units, with PDMS top layer and air layer dimensions set at 50 μm. Numerical simulation indicated that acoustic pressure nodes (i.e., pressure zero regions) overlaid with the design of air-filling units when the acoustic wavelength was ten times or more than the thickness of both PDMS top and air layers (f =2 MHz). When the acoustic wavelength was comparable with either PDMS top or air layer thickness, the pressure nodes didn’t overlay the predesigned air-filling units (Figure 2A). 3D modeling and simulation further illustrated that the air-filling units were pressure zero, resulting in near-field acoustic pressure well existing above the air-filling unit on the surface of the PDMS top layer (Fig. 2B). Additionally, numerical simulation of acoustofluidic velocity field demonstrated the velocity zero regions distributed around the edge of the air-filling unit (Fig. 2C), indicating the microparticle tends to concentrate around the edge of the unit. We observed that the simulated pressure nodes overlaid with microparticle assembly pattern (Figure 2D). Furthermore, quantitative image analysis also indicated pressure wells were highly consistent particle aggregation regions (Figure 2E), confirming that SLAB belonged to acoustic node bioassembly.

**Figure 2:**
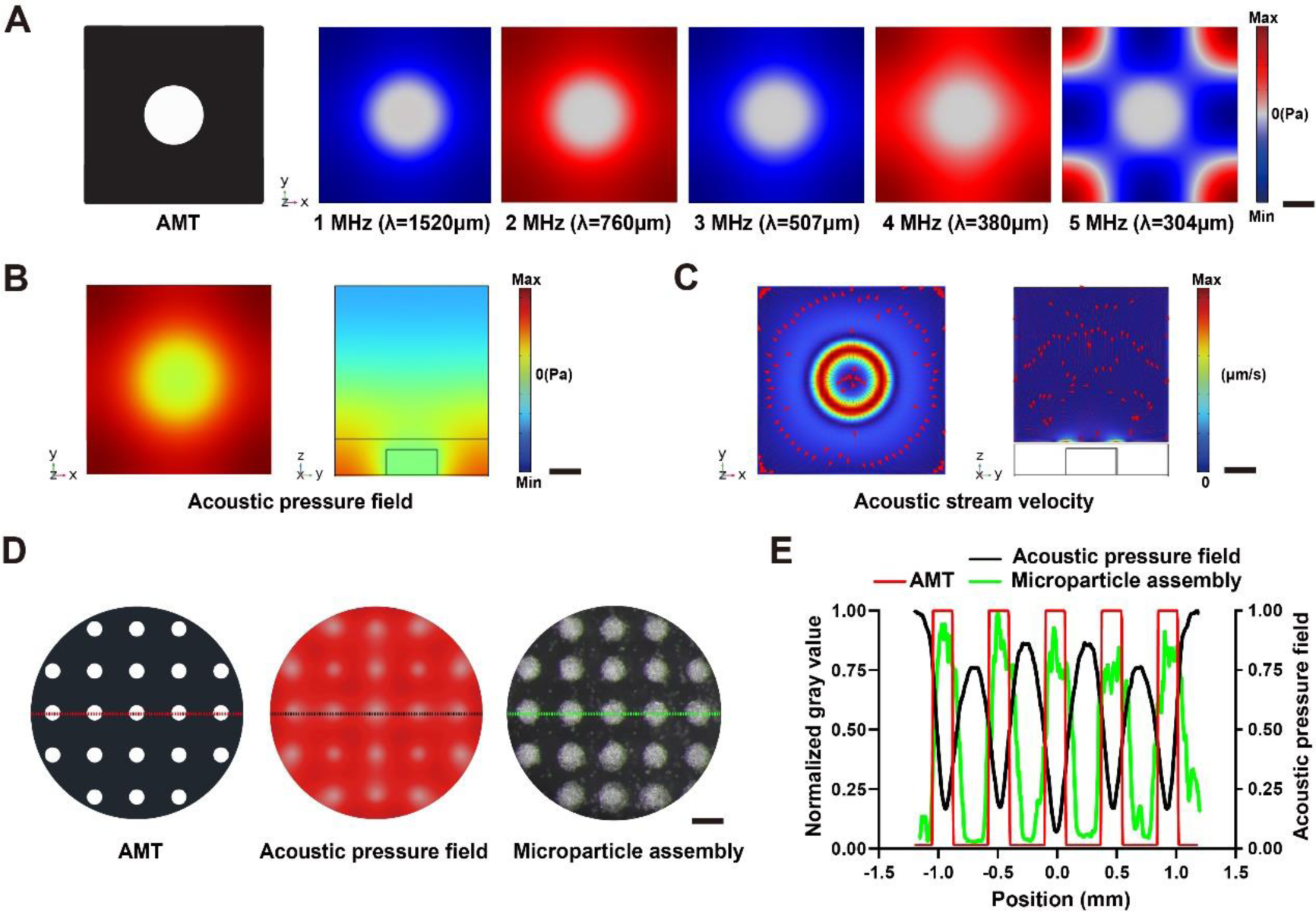
Numerical simulation of acoustic waves. (A) Acoustic pressure field in the XY plane of the circular pattern under varied excitation frequencies (1, 2, 3, 4, and 5 MHz). (B) Acoustic pressure fields in the XY and XZ planes of the circular pattern shaped air-filling region. (C) Acoustic stream velocity in the XY and XZ planes of the circular pattern shaped air-filling region. (D) The target pattern of AMT and corresponding acoustic pressure field and microparticle assembly. (E) Quantitative comparison of AMT and corresponding acoustic pressure field and microparticle assembly. Scale bar: (A), (B) &(C) 100 μm, (D) 200 μm.

### 3.2. Microparticle assembly experiments

The simulation results were validated through microparticle assembly experiments performed after optimizing the PDMS top layer and air layer of the AMT. We examined the assembly capability of SLAB for microparticles with various diameters. Specifically, SLAB was utilized to assemble PS microparticles with diameters of 10, 50, 100, and 200 μm onto a target pattern of a 200 μm diameter circle (Fig. S1A and Video S1). The result demonstrated that microparticles were efficiently assembled on the top of the target pattern when the microparticle size was smaller than the target pattern size. Additionally, microparticle assembly experiments demonstrated that initially randomly distributed PS microparticles aggregated into a circular array within 60 seconds, leaving space in the antinode area. Furthermore, we identified that a minimal concentration of 1.5 mg/mL microparticles was requested to fully overlay the target pattern when the microparticle size was 10 μm (Fig. S1B-C). We also investigated the size effect of circular array patterns on the assembly (Fig. S1D). The numerical simulation depicted that pressure node regions overlaid with target circular patterns with varied diameters (50, 100, and 200 μm). Microparticle assembly experiments further confirmed that the microparticles with the size of 10 μm were assembled on the top of the target patterns regardless of the pattern size.

Symmetrical and asymmetrical human bionic structures were designed to evaluate the versatility of SLAB in constructing arbitrary target patterns, including the hepatic lobules, muscle fibers, osteons, and microvessels (Fig. 3). The numerical simulations indicated that acoustic pressure node regions fitted well with the target patterns regardless of symmetrical or asymmetrical (Fig. 3A-B). Particle tracking analysis further demonstrated that initially randomly distributed microparticles on the AMT were rapidly trapped in the acoustic pressure node regions, forming tightly arranged particle aggregates (Video S2). Microparticle assembly experiments further verified that 91.6% of microparticles were assembled onto the target pattern regions (Video S3).

**Figure 3:**
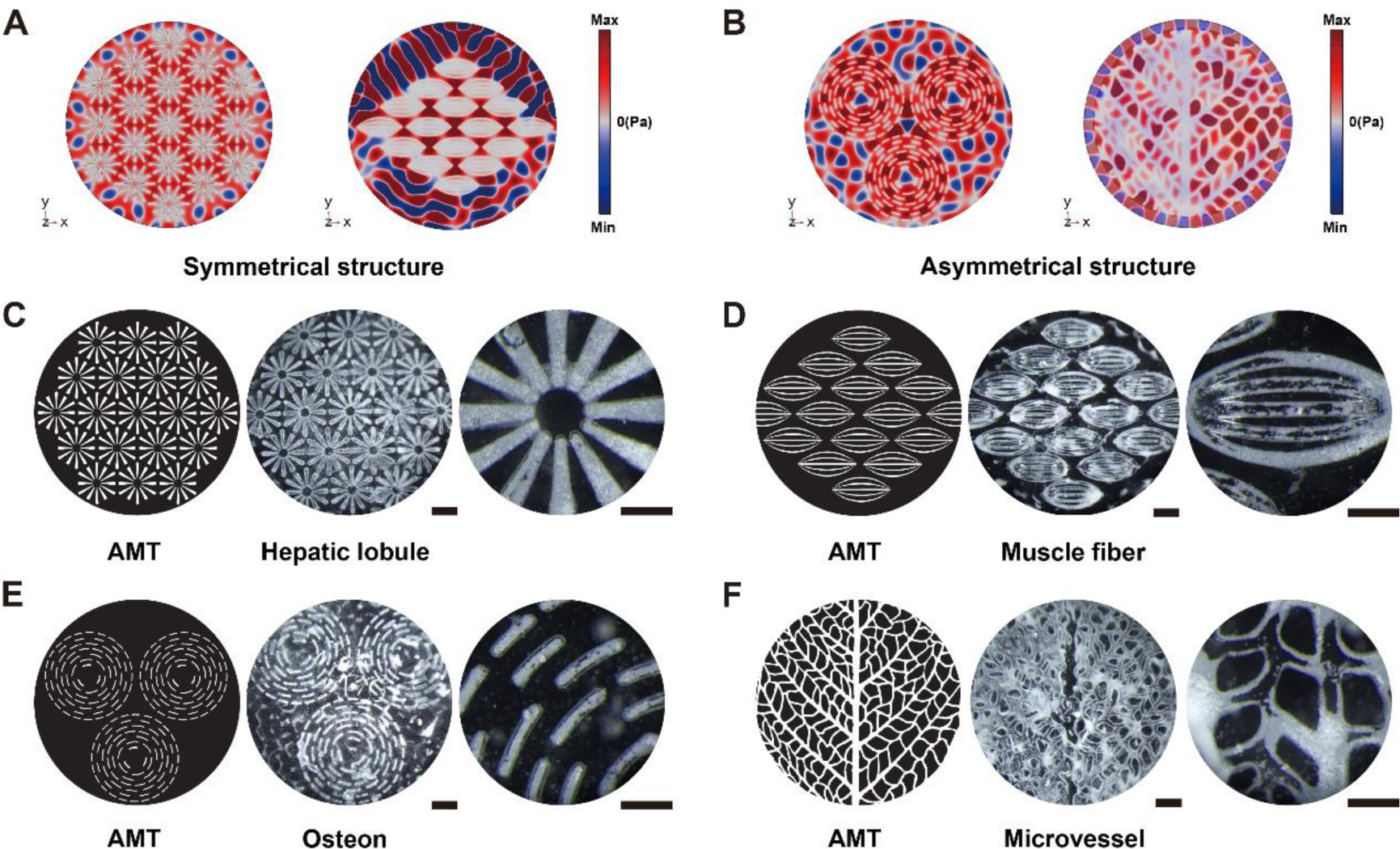
Microparticle assembly of symmetrical and asymmetrical bionic structure. (A) Numerical simulation of acoustic pressure fields formed by target patterns with symmetrical bionic structures. (B) Numerical simulation of acoustic pressure fields formed by target patterns with asymmetrical bionic structures. (C) Microparticle assembly of hepatic lobule pattern. (D) Microparticle assembly of muscle fiber pattern. (E) Microparticle assembly of osteon pattern. (F) Microparticle assembly of microvessel pattern. Scale bars: (C), (D), (E) &(F) 1 mm, 500 μm.

### 3.3. Construction of LTMs using SLAB

In the liver tissues, the hepatic lobule serves as the fundamental functional unit, featuring hepatic cell cords and hepatic blood sinuses arranged radially around the central vein, creating a hexagonal heterogeneous cell structure (Fig. 4A). The LTMs were generated through the acoustic assembly of hepatic cells and endothelial spheroids, followed by a 7-day tissue culture (Fig. 4B). Endothelial spheroids were generated in the ultra-low adhesion agarose microporous arrays with dynamic culture on a shaker for 4 days (Fig. S2A). The average diameter of the spheroids reached 143 ± 5.83 µm (Fig. S2B). Endothelial spheroids were assembled in the radial area of the target pattern, and hepatic cells were evenly dispersed throughout the bioassembly chamber. The biocompatibility of the SLAB was assessed through cell viability assays. Dead cells exhibited red fluorescence through PI staining, while total nuclei emitted blue fluorescence via Hoechst 33342 staining (Fig. 4C). No significant difference in cell viability was observed before and after assembly, indicating that the assembly process and subsequent culture did not affect cell viability (Fig. 4D).

**Figure 4:**
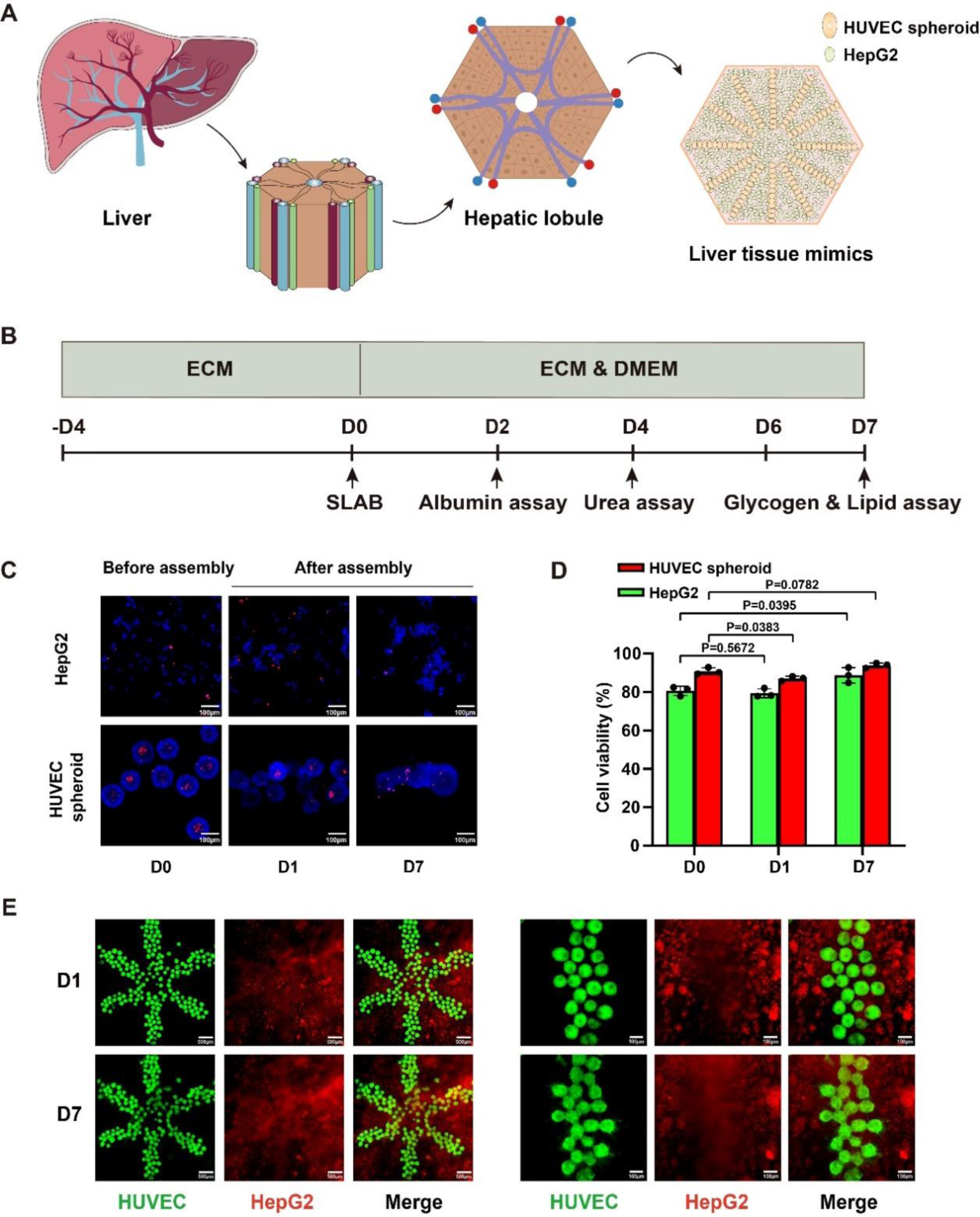
Construction of LTMs using SLAB. (A) Schematic illustration of cytoarchitecture of a hepatic lobule. (B) Experimental designs of the LTM production and characterization. (C) Evaluation of cell viability of the SLAB and control groups at day 0, day 1 and day 7. (D) Quantitative analysis of cell viability of the SLAB and control groups. (E) The fluorescence image of the LTMs at day 1 and day 7. Scale bars: (C) 100 μm, (E) 500 μm, 100 μm.

Furthermore, we performed cell tracking analysis on the LTMs after assembly and after 7 days of culture. The results showed that endothelial spheroids were assembled into the radial area of the hepatic lobular pattern while hepatic cells tightly surrounded the endothelial cells and filled the rest areas of the assembly chamber (Fig. S3A and S3B). Thus, LTMs emulated the natural hepatic lobule structure and enabled direct contact between liver parenchyma cells and supporting cells (Fig. 4E and Video S4). The LTMs merged into a piece of tissue within 7 days without losing predefined heterotypic cell structure.

### 3.4. Functional evaluation of the LTMs

We detected albumin secretion and urea production on day 2, 4, and 6 to evaluate the liver metabolism and synthesis functions of the LTMs. The albumin secretion and urea production of the LTMs reached the highest level on day 6. Compared to the non-assembed group, the SLAB group displays improved secretion functions in albumin secretion (17.34 ± 0.27 vs. 14.46 ± 0.21 ng/mL) (Fig. 5A) and urea secretion (1.62 ± 0.06 vs. 1.19 ± 0.04 mg/dL) (Fig. 5B) on day 6. In addition, PAS and Nile red staining showed that the LTMs had liver-specific functions, including glycogen storage and lipid accumulation (Fig. 5C and 5D). The SLAB group showed significantly larger lipid droplets (104% increase in diameter and 651% increase in volume) than the non-assembled group (Fig. 5E).

**Figure 5:**
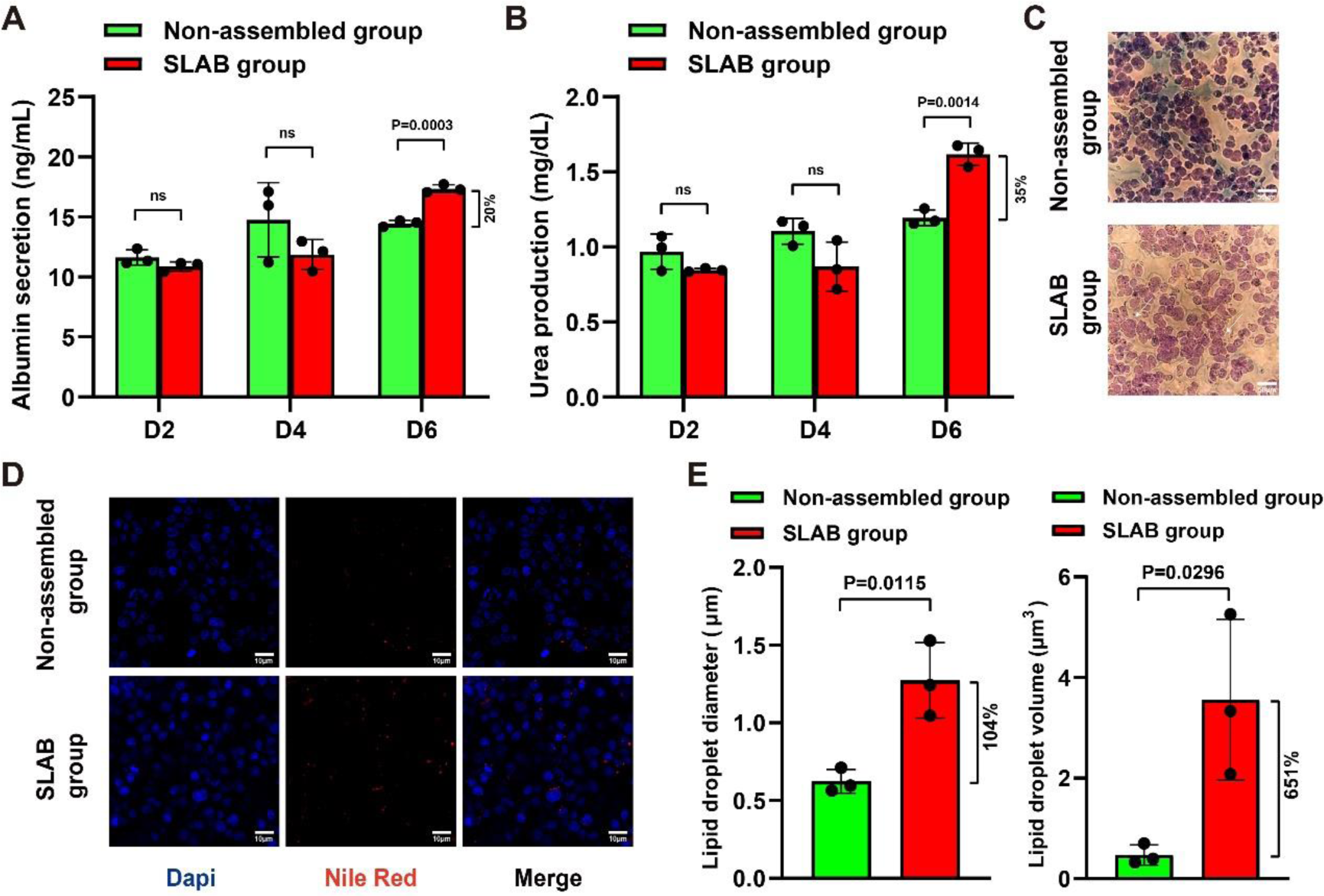
Liver function assays for the LTMs. (A) Analysis of albumin secretion by ELISA. (B) Analysis of urea production by quantitative colorimetric assay. (C) Glycogen storage assessment by Periodic-Acid Schiff (PAS) staining. (D) Lipid accumulation assessment by Nile red staining (E) Quantitative analysis of lipid production. Scale bars: (C) 50 μm, (D) 10 μm.

## 4. Discussion

The capacity to generate arbitrarily shaped multicellular structures is an essential basis for advanced biofabrication and their application in constructing complex tissue mimics. Although acoustic bioassembly has demonstrated great promise to become an efficient biofabrication tool to generate functional tissue mimics, most acoustic bioassembly techniques are currently limited to generate acoustic node or antinode patterns with periodic and symmetrical structures. Here, we have demonstrated the SLAB technique as a novel acoustic bioassembly technique to generate geometrically defined arbitrary multicellular structures. SLAB employs amplitude modulation of acoustic traveling waves and generates arbitrary acoustic pressure distribution via an AMT. Specifically, the AMT is based on a microfabricated PDMS construct that is fabricated using soft lithography. Soft lithography is a widely accessible microfabrication technique, and it’s broadly utilized for manufacturing microfluidic chips. Soft lithography enables to fabricate a feature structure down to 500 nm. Experimentally, we have demonstrated the acoustic bioassembly of microparticles using a PDMS-based acoustic template with a feature size down to 25 μm. Additionally, we have examined the geometric effects of the PDMS construct on acoustic bioassembly and found that the PDMS top layer and air layer thickness play a critical role in modulating acoustic pressure distribution. Particularly, when both the PDMS top layer and air layer thickness are approximately one-tenth of the wavelength, the air-filled region works as a sound barrier and prevents incident waves from passing through the PDMS construct. Thus, the spatial distribution of zero pressure areas of the near-field acoustic waves at the surface of PDMS layer overlays with the air layer of PDMS construct. Using this technique, bioparticles, such as cell aggregates with diameters ranging from 10 to 200 μm, can be assembled into arbitrarily-geometric structures predefined by soft-lithography-defined template.

SLAB exhibits several unique advantages in generating geometrically defined arbitrary multicellular structures over the existing acoustic bioassembly techniques. Specifically, standing wave based acoustic bioassembly techniques have been demonstrated to construct various types of tissue mimics, including endothelium, brain tissue, and heart tissue^31–34^. However, the bioassembly patterns of these tissue mimics are limited to rings, hexagons, stripes, grids, and crystal-like symmetrical and periodic structures^35–39^. Despite emerging acoustic phase holography techniques permitting the formation of arbitrarily-geometric patterns, these techniques highly rely on complex algorithms and high-precision 3D printing to design and fabricate acoustic metamaterial for phase modulation^40–44^. Distinct from these phase holography techniques, SLAB directly modulates the distribution of acoustic pressure field using air regions of microfabricated PDMS construct, eliminating complex calculation and high-precision fabrication of phase modulation metamaterial. More importantly, SLAB theoretically offers a spatial resolution equal to traditional UV photolithography. In contrast, the spatial resolution of phase holography techniques is determined by acoustic holographic kinoform fabricated by 3D printing techniques. Thus, the feature width of the assembled structure is limited to tens of micrometers. Additionally, SLAB allows the formation of multicellular structure at the bottom of a liquid layer, thus providing a more biocompatible environment to most cell types than the acoustic bioassembly techniques that request cells at the air-liquid interface^45–50^.

We employ SLAB to construct LTMs. Specifically, we emulate radially-like hepatic sinusoids around the central vein of the hepatic lobules in the native liver tissues and design a PDMS construct with radially-like air-filling regions. By exploring the different acoustic impedances of endothelial spheroids and hepatic cells, we assemble endothelial spheroids into predefined radially-like arrangement surrounded by hepatic cells. We analyze a functional assessment of the assembled LTMs during a 7-day tissue culture. Compared to the non-assembed group, the SLAB group displays improved secretion function in albumin secretion (17.34 ± 0.27 vs. 14.46 ± 0.21 ng/mL) and urea secretion (1.62 ± 0.06 vs. 1.19 ± 0.04 mg/dL) on day 6. Additionally, the SLAB group demonstrates enhanced functions of glucolipid metabolism in terms of lipid storage and glycogen accumulation compared to the non-assembled group. Specifically, the size of lipid droplets in the SLAB group is significantly larger than that in the non-assembled group. Previously, Faraday wave differential bioassembly has been employed to generate hepatic lobule-like tissue mimics from hepatic spheroids and endothelial encapsulating microgels^51^. However, Faraday wave bioassembly is more suitable to generate centimeter-scale tissue mimics with a feature dimension of millimeter-scale due to their low working frequencies (<1000 Hz). In contrast, SLAB enables to generate millimeter-scale tissue mimics with a feature dimension of tens of micrometers. Thus, SLAB could provide a more precise emulation of the hepatic functional units. In addition, 3D bioprinting has been demonstrated to construct hepatic-lobule like tissue mimics using digital micromirror device (DMD) system^52, 53^. However, the live cells within the printed biomaterials are randomly distributed and lack close intercellular contact. In contrast, SLAB utilizes physical field interactions with biological elements to directly manipulate cells. Cells self-assemble into densely packed structures where force potential is lowest, promoting cell communication and enhancing tissue-specific functions.

## 5. Conclusions

Overall, we have demonstrated SLAB, a novel acoustic bioassembly technique, to generate geometrically defined arbitrary multicellular constructs. The SLAB explores widely accessible soft lithography and enables to generate asymmetrical heterotypic patterns. The SLAB procedure is facile and biofriendly with live cells. We anticipate this bioassembly technique will serve as a critical tool to promote applications of acoustic bioassembly in tissue engineering and regenerative medicine.

## Author contributions

Conceptualization: P.C..

Data curation: S.C., P.C..

Formal analysis: S.C., P.C..

Funding acquisition: P.C..

Investigation: S.C., J.W., Q.J., P.C..

Methodology: S.C., P.C..

Project administration: P.C..

Resources: P.C, Q.Z..

Software: S.C., J.W., Y.F..

Supervision: P.C., Q.Z., Y.C..

Validation: P.C., J.W..

Visualization: S.C., Y.Z., J.W..

Writing—original draft: S.C., Z.W., P.C..

Writing—review & editing: P.C., W.Z., Y.C..

## Data and materials availability

All data needed to evaluate the conclusions are present in the paper and the Supplementary Materials. Additional data related to this paper may be requested from the authors.

## Supplementary Figures

**Figure S1:**
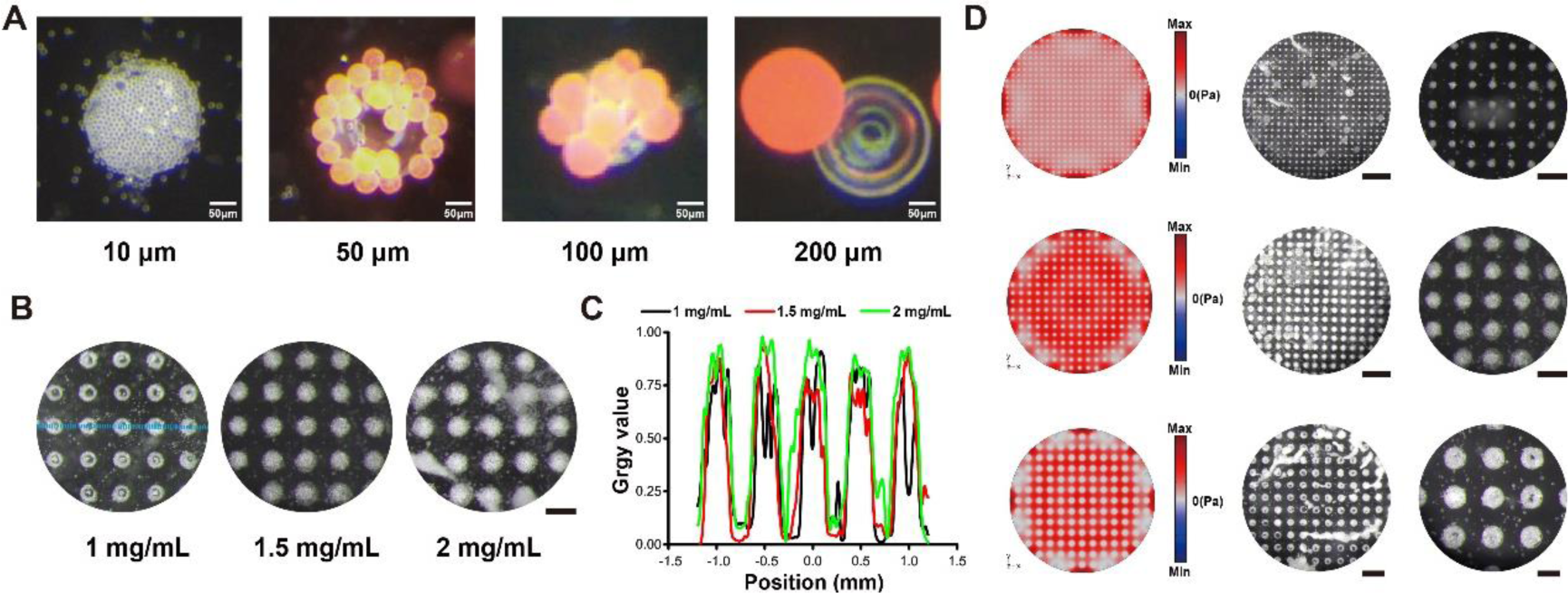
Soft-lithographically defined acoustic assembly of microparticles. (A) The PS microparticles with different diameters (10, 50, 100, and 200 μm) were assembled on the target circular patterns with a size of 200 μm. (B) Microparticle assembly under varied microparticle concentrations (1, 1.5, and 2 mg/mL). (C) Quantitative analysis of microparticle assembly under varied microparticle concentrations. (D) Numerical simulations of acoustic pressure field and corresponding microparticle assembly (10 μm) into circular target patterns with varied diameters (50, 100, and 200 μm). Scale bars: (A) 500 μm, (B) 200 μm, (D) 1 mm, 500 μm.

**Figure S2:**
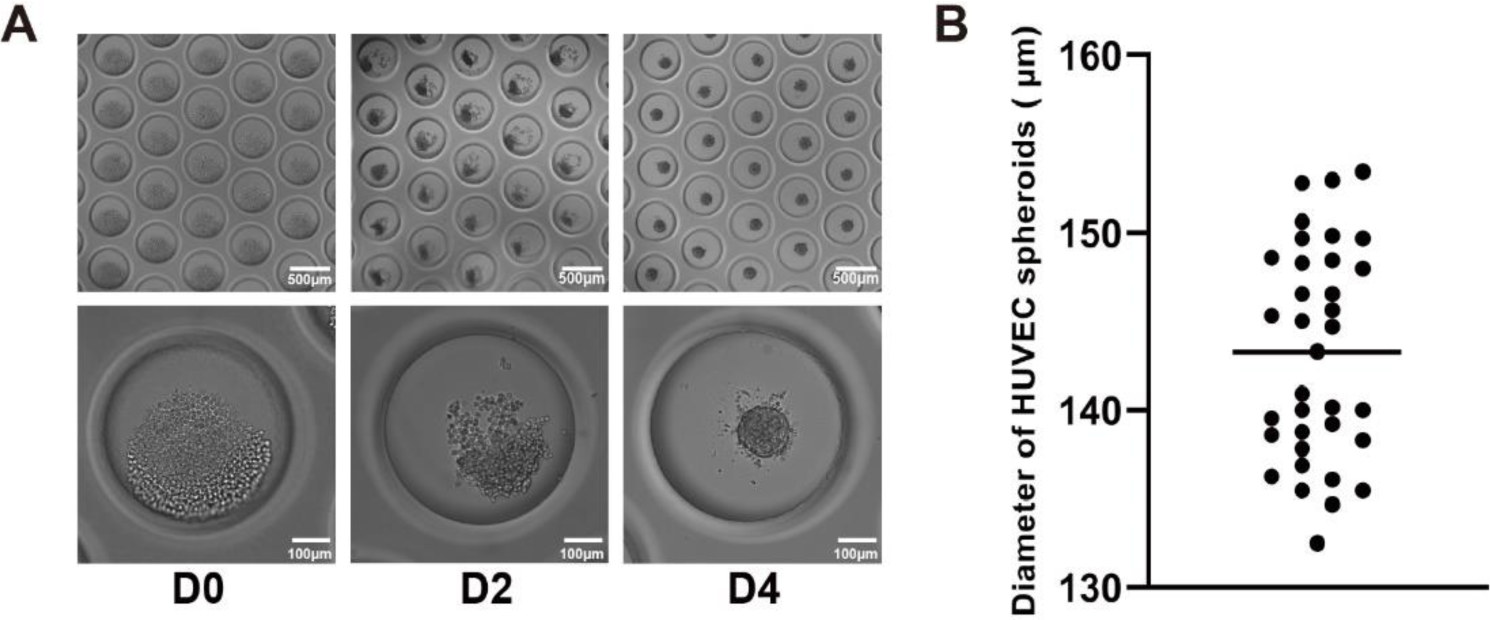
Generation of endothelial spheroids. (A) The bright-field images of endothelial spheroids formed in an agarose microcavity array. (B) Size distribution of endothelial spheroids. Scale bars: (A) 500 μm, 100 μm.

**Figure S3:**
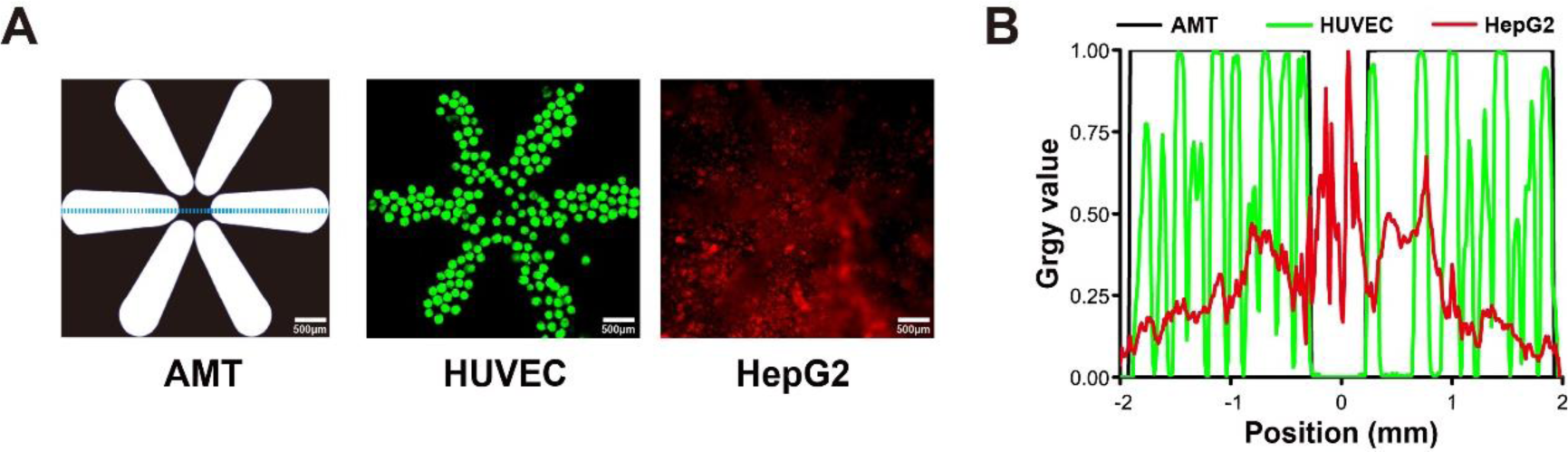
Distribution analysis of LTMs. (A) The target pattern unit and the corresponding fluorescence images of assembled HUVEC spheroids and HepG2 cells. (B) Distribution analysis results along the XY plane of HUVEC and HepG2 cells. Scale bar: (A) 500 μm.

## Supplementary Tables

**Table 1.** The regent information.

| Parameter | Symbol | Value |
| --- | --- | --- |
| Speed of sound in air | $c_1$ | $343 \text{ m}\cdot\text{s}^{-1}$ |
| Speed of sound in PDMS | $c_2$ | $1076.5 \text{ m}\cdot\text{s}^{-1}$ |
| Speed of sound in water | $c_3$ | $1520 \text{ m}\cdot\text{s}^{-1}$ |
| Viscosity | $\eta$ | $0.0025 \text{ Pa}\cdot\text{s}$ |
| Density | $\rho$ | $1020 \text{ Kg}\cdot\text{m}^{-3}$ |
| Frequency | $f$ | $1.9 \text{ MHz}$ |
| Electric potential | $V_{pp}$ | $10 \text{ V}$ |
| Radius of particle | $r$ | $10 \text{ }\mu\text{m}$ |

**Table 2.**
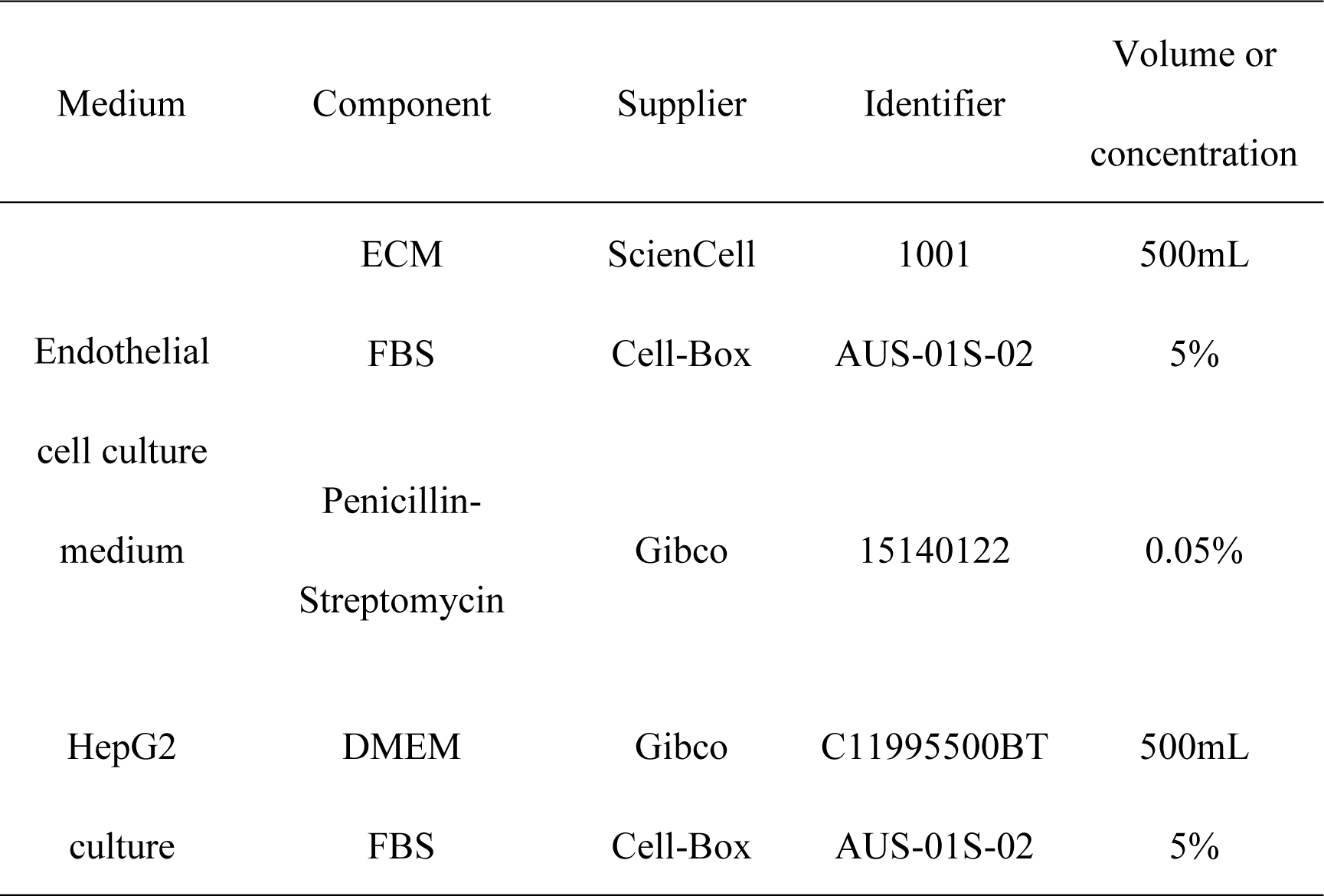
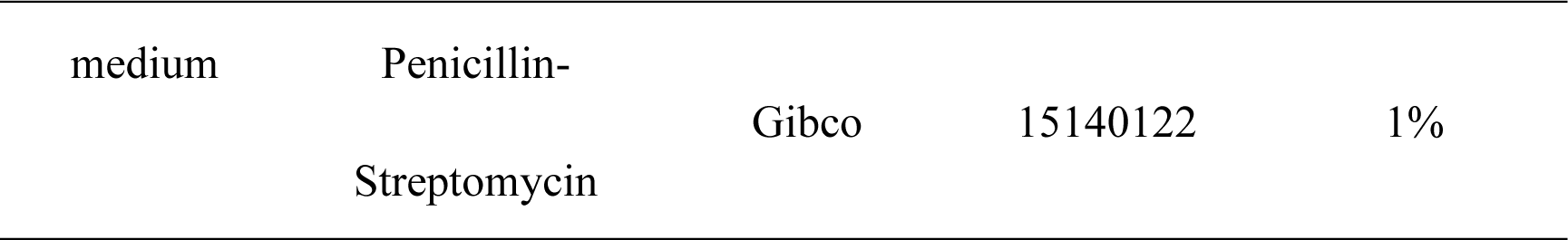
The parameters of reagents.

